# Development of affinity beads-based *in vitro* metal-ligand binding assay reveals dominant cadmium affinity of thiol-rich small peptides phytochelatins beyond glutathione

**DOI:** 10.1101/2021.08.23.456032

**Authors:** Shimpei Uraguchi, Kenichiro Nagai, Fumii Naruse, Yuto Otsuka, Yuka Ohshiro, Ryosuke Nakamura, Yasukazu Takanezawa, Masako Kiyono

## Abstract

For a better understanding of metal-ligand interaction and its function in cells, we developed an easy, sensitive, and high-throughput method to quantify ligand-metal(loid) binding affinity under physiological conditions by combining ligand-attached affinity beads and inductively coupled plasma-optical emission spectrometry (ICP-OES). Glutathione (GSH) and two phytochelatins (PC2 and PC3, small peptides with different numbers of free thiols) were employed as model ligands and attached to hydrophilic beads. The principle of the assay resembles that of affinity purification of proteins in biochemistry: metals binding to the ligand on the beads and the rest in the buffer are separated by a spin-column and quantified by ICP-OES. The binding assay using the GSH-attached beads and various metal(loid)s suggested the different affinity of the metal-GSH interactions, in accordance with the order of the Irving–Williams series and the reported stability constants. The binding assay using PC2 or PC3-attached beads suggested positive binding between PCs and Ni(II), Cu(II), Zn(II), Cd(II), and As(III) in accordance with the number of thiols in PC2 and PC3. We then conducted the competition assay using Cd(II), Mn(II), Fe(II), Cu(II), and Zn(II) and the results suggested a better binding affinity of PC2 with Cd(II) than with the essential metals. Another competition assay using PC2 and GSH suggested a robust binding affinity between PCs and Cd(II) compared to GSH and Cd(II). These results suggested the dominance of PC-Cd complex formation *in vitro*, supporting the physiological importance of PCs for the detoxification of cadmium *in vivo*. We also discuss the potential application of the assay.

## Introduction

Essential metals are utilized by cells as cofactors for many proteins [1]. A bioinformatics study estimated that approximately 40% of the enzymes with known three-dimensional structures required metals at their catalytic centers [2]. The proportions of essential metals associated with enzymes are 6–9% of the analyzed enzymes for zinc (Zn), iron (Fe), and manganese (Mn), respectively, and 1% for copper (Cu). These metals are preferentially utilized in respective Enzyme Commission (EC) classes: for instance, Fe is mostly utilized by oxidoreductases (EC1) and Zn-containing enzymes are prominently enriched in hydrolases (EC3)[2]. Zn also plays structural roles in Zn-binding proteins containing zinc-finger domains, suggesting the importance of Zn in transcriptional and post-translational regulations, especially in eukaryotes [3,4].

Meanwhile, the essential heavy metals in the cells are tightly regulated to avoid their potential toxicity. The studies on bacteria and yeast suggest that there is practically no cytosolic pool of free Cu and Zn ions under steady-state conditions [5–7]. The essential heavy metals imported to the cells are preferentially delivered to the target proteins requiring respective metals, and the rest are “buffered”, bound to various small-molecular ligands as well as proteins like metallothionein in mammals [8,9]. Under conditions with excess metals, the biological chelators then function in “muffling”, chelating the excess metal ions in the cell and avoiding cytotoxicity [8,9].

Among such metabolites, GSH is believed to be a major metal-chelating small molecule especially in mammalian cells [8,10]. In addition, many diverse small-molecular ligands are physiologically significant metal chelators in bacteria, yeast, worm, and higher plants [11–13]. (Phyto)siderophores and its precursor nicotianamine (NA) are typical examples that function as Fe(II) and Zn(II) binding ligands, respectively. Other established examples of the small molecular chelators are phytochelatins (PCs), cysteine-rich small peptides. The general structure of PCs is (γ-Glu-Cys)n-Gly with *n* usually between 2 and 7 [14]. PCs are non-ribosomally synthesized from GSH (γ-Glu-Cys-Gly) by phytochelatin synthases (PCSs). Compared to (phyto)siderophores and NA that function as ligands for essential heavy metals [12], PCs can be characterized as high-affinity chelators for a wide range of toxic metal(loid)s such as cadmium (Cd) and arsenic (As). In plants, PCs play a central role in the detoxification of various toxic metal(loid)s [15–19]. Interestingly, the PC-mediated detoxification of toxic metal(loid)s is also demonstrated in the fission yeast *Schizosaccharomyces pombe* [20], in which PCs were first discovered [21], and in the nematode worm *Caenorhabditis elegans* [22]. These suggest the conserved functions of PC/PCS for toxic metal(loid) detoxification in taxonomically distant organisms.

The physiological significance of GSH and PCs in metal(loid) detoxification is mostly demonstrated by phenotypes induced by GSH synthesis inhibitors or the disruption of the genes for PC/GSH synthesis, which lead to metal(loid) hypersensitivity [15–17,22,23]. These results indicate that ligand-metal complex formation is a key process for the PC/GSH-mediated toxic metal(loid) detoxification, however, there is limited information on complex formation between PCs/GSH and metal(loid)s, especially under physiologically relevant conditions. It appears first due to the technical difficulties overall [24], and secondly, due to unfamiliarity with required analytical techniques and/or limited access to the analytical devices especially for many molecular metal-biologists. For example, the PCs/GSH-metal(loid) complex formation *in vitro* is often examined by potentiometry, voltammetry, and nuclear magnetic resonance (NMR) spectrometry [25]. Isothermal titration calorimetry (ITC) is also a promising approach, providing detailed *in vitro* data of binding chemistry under physiological conditions, but the device is expensive and not widely available. Another disadvantage of these analytical chemistry approaches for molecular metal-biologists is that it is technically difficult to mimic a cellular matrix containing various metals and metabolites other than the targets, which may also be potential interactants. Liquid-chromatography-inductively coupled plasma mass-spectrometry (LC-ICP-MS) and liquid-chromatography-electrospray ionization mass-spectrometry (LC-ESI-MS) enable direct detection of multiple metal-ligand complexes *in vitro* and *in vivo*, however, the technical difficulty of analyzing the intact complex remains the bottleneck [26].

For these reasons, an alternative assay especially for metal biologists would be a useful option to advance our understanding of metal-ligand interactions at conditions that are prevalent in the cytosol. The present study aimed to develop an ICP-based simple but quantitative method to evaluate ligand-metal(loid) interaction under various physiological conditions. ICP-optical emission spectrometry (OES) was used as an analytical tool. ICP-OES as well as ICP-MS is a common device among metal-biologists, compared to the other analytical techniques, and more importantly, ICP-based element analysis provides highly sensitive and high-throughput quantification. We employed GSH and PCs as model ligands and prepared ligand-attached affinity beads for the assay. The GSH-metal(loid)s binding ratios derived from our method coincided with the reported values. With the established assay, we examined PC and Cd(II) interactions under physiological conditions containing GSH and several essential metals. The competitive assays demonstrated preferential interaction of PCs and Cd(II), which supports the *in vivo* observations that PCs play a central role in Cd detoxification in plants, fission yeast, and *C. elegans*.

## Materials and methods

### Synthesis of PC2 and PC3

PC2 and PC3 were synthesized by the Fmoc solid-phase peptide synthesis strategy [27] and the purity and mass sizes were verified by LC-UV/MS analysis. A full method was provided as supplemental materials and methods (see Supplementary information). LC-MS was performed on an Agilent1200 system (G1379B degasser, G1312A binary pump, G1329A auto liquid sampler, G1316A thermostat column compartment, and G1315B diode array detector) and a JEOL AccuTOF LC-plus T100LP time-of-flight (TOF) mass spectrometer equipped with an ESI interface. The acquisition mass range was *m/z* 100–1000. Mobile phase A was 0.1% HCO_2_H in H_2_O, and mobile phase B was 0.1% HCO_2_H in CH_3_CN. Separations on the LC-MS were performed on a CAPCELL CORE C18 (2.7 μm, 2.1 × 50 mm, SHISEIDO Co., Ltd., Tokyo, Japan) using a linear gradient of 5–100% solvent B over 8 min at a flow rate of 0.4 mL/min. Preparative HPLC was performed on a GL sciences system (GL7420 autosampler, GL7410 HPLC pump, and GL7450 UV detector). Separations on the HPLC were performed on a CAPCELL PAK C18 (Type: UG120, 5 μm, 20 × 250 mm, SHISEIDO Co., Ltd., Tokyo, Japan) using 5% mobile phase B was used at a flow rate of 8 mL/min. UV detection was at 210 nm. UV–vis spectra were obtained on a HITACHI U-2800 UV spectrophotometer with 1 cm quartz cells.

### Loading of GSH, PC2, or PC3 onto the affinity beads

TOYOPEARL AF-Formyl-650 (TOSOH, Tokyo, Japan) was selected as the basal beads. TOYOPEARL AF-Formyl-650 (1 mL, 60 μmol/mL) was placed in a polypropylene tube and washed with H_2_O (1 mL, 1 min × 3). To the beads was added a solution of GSH (18 mg, 60.0 μmol) in 0.1 M phosphate buffer (pH 7.0, 1 mL), and then sodium cyanoborohydride (11 mg, 180 μmol) was added. The resulting mixture was agitated at room temperature for 22 h. The solvents and soluble reagents were removed by filtration, and the bead was washed with H_2_O (1 mL, 1 min × 3), saturated sodium chloride (1 mL, 1 min × 3), H_2_O (1 mL, 1 min × 3), and MeOH (1 mL, 1 min × 3). PC2 or PC3 were loaded onto the affinity bead according to the loading procedure described above. The loading of the resulting bead was determined by the amount of 2-nitro-5-thiobenzoate anion formed from the reaction of 5,5’-dithiobis(2-nitrobenzoic acid), Ellman’s reagent, with sulfhydryl groups on the bead. The affinity resin (10 mg) was treated with 2.5 mM Ellman’s reagent in MeOH (2 mL) and *N*,*N*-diisopropylethylamine (5 µL) at room temperature for 30 min. The total volume was adjusted to 5 mL by the addition of MeOH and measured spectrophotometrically at 412nm.

### Single metal(loid)-beads assay

The interaction assay was conducted in a 5 mL Eppendorf tube containing 3.0 mg of the beads (control, GSH, PC2, or PC3). The assay was repeated with at least three replications. To evaluate the interaction between the ligand beads and various metal(loid)s, 5 mL of 40 mM HEPES (pH 7.2) containing 200 nmol of MnCl_2_, FeCl_2_, (NH_4_)_2_Ni(SO_4_)_2_, CuSO_4_, ZnSO_4_, NaAsO_2_, or CdCl_2_, was added to the 5 mL tube. The tube was immediately set to a rotator (RT-50, TAITEC, Saitama, Japan) and incubated at room temperature for 5 min at approximately 4 rpm. For Mn(II), Fe(II), Cu(II), and As(III), overnight incubation was also tested with rotation. The metal reagents used in this study were all Guaranteed Reagents grade purchased from Nacalai Tesque (Kyoto, Japan) or FUJIFILM Wako Pure Chemical (Osaka, Japan). The reagent for NaAsO_2_ was USP grade, purchased from Merck (Darmstadt, Germany).

### Multiple metals competition assay

The competition assay using multiple metals was conducted as described for the single metal(loid)-beads assay, except for the metal composition in the reaction buffer. The reaction buffer [5 mL of 40 mM HEPES (pH 7.2)] contained 20 nmol of MnCl_2_, 40 nmol of FeCl_2_, 2 nmol of CuSO_4_, 20 nmol of ZnSO_4_, and 200 nmol of CdCl_2_ and was added to 3.0 mg of the beads (control, GSH, PC2, or PC3) in the 5 mL tube. The tube was immediately set to a rotator and incubated at room temperature for 5 min. The assay was repeated with three independent reactions.

### PC2 and GSH competition assay

Cd-binding competition between PC2 and GSH was examined by modifying the single metal(loid)-beads assay. For the assay using GSH-attached beads (3.0 mg), the reaction buffer containing 40 mM HEPES (pH 7.2), 100 nmol of CdCl_2_, and 100 nmol of PC2 was used. For the assay using PC2-attached beads (3.0 mg), the reaction buffer containing 40 mM HEPES (pH 7.2), 100 nmol of CdCl_2_, and 25 or 200 nmol of GSH was used. The prepared tube was immediately set to a rotator and incubated at room temperature for 5 min. The assay was repeated with three independent reactions.

### Quantification of metal(loid) binding to the ligand

For Protocol A (Fig. 2), which was used for most of the assay in this study, an aliquot (720 µL) of the reaction mixture was immediately applied to an empty spin column (EconoSpin, Ajinomoto Bio-Pharma, Osaka, Japan) which harbors a polyethylene membrane, placed in a 1.5 mL plastic tube. The affinity beads were removed from the mixture by immediately centrifuging the column at 15,300 g for 10 sec using a standard microcentrifuge. An aliquot (500 µL) of the flow-through was added to 500 µL of 60% nitric acid (for poisonous metals analysis, FUJIFILM Wako Pure Chemical, Osaka, Japan) and then diluted to 5 mL with MilliQ water. Elemental concentrations in the diluted samples were determined by ICP-OES (iCAP7400Duo, Thermo-Fisher Scientific, Waltham, MA). Emission wavelengths used for the detection were 189.04, 206.200, 224.700, 226.502, 239.562, and 260.569 nm for As, Zn, Cu, Cd, Fe, and Mn, respectively. Then the residual (free) metal(loid) quantity in the reaction mixture after the reaction was calculated for control beads (M_[control filtrate]_) and ligand-attached beads (M_[ligand filtrate]_), respectively (Fig. 2). The amount of the metal(loid) that interacted with the ligand attached to the beads (M_[binding]_) was obtained by subtracting M_[ligand filtrate]_ from M_[control filtrate]_ (Fig. 2). The metal (loid)/ligand binding ratio was calculated by dividing M_[binding]_ by the ligand amount in the reaction mixture.

**Fig. 2.**
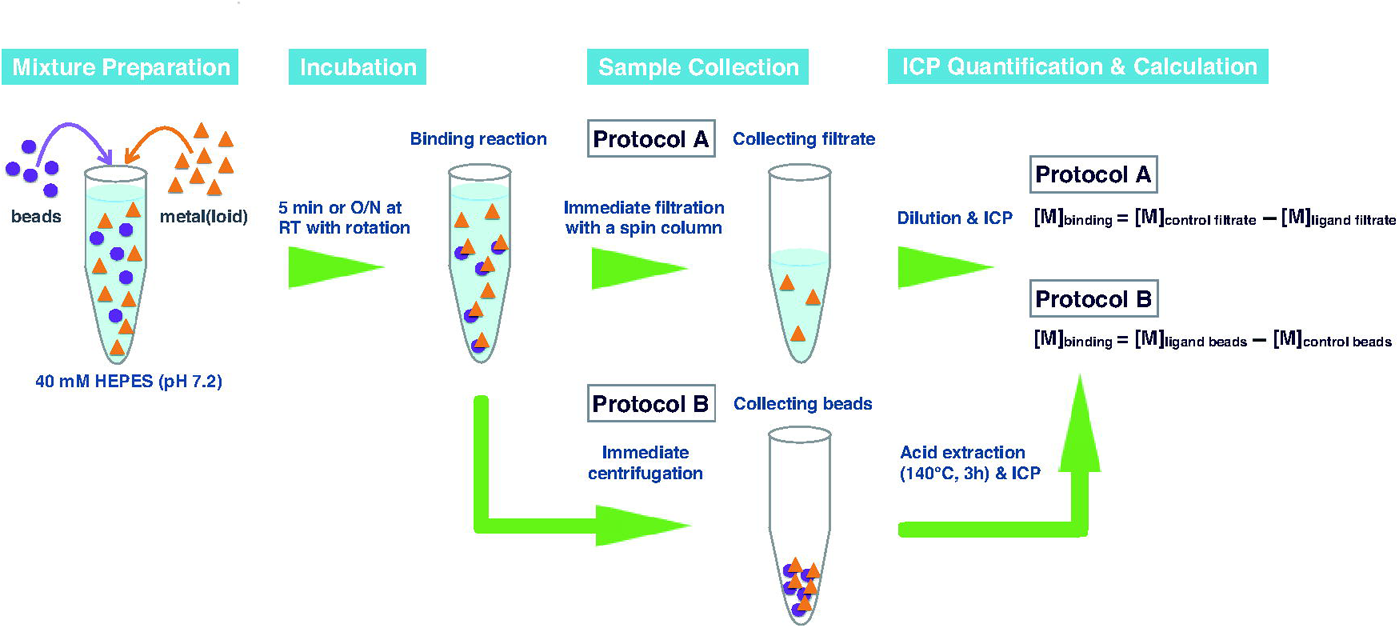
Schematic diagram describing the two protocols of the metal-ligand binding assay using the ligand-attached beads and ICP. A reaction mixture for the standard protocol was prepared in a 5 ml plastic tube containing the beads (normally 3.0 mg) and a metal(loid)-of-interest (normally 200 nmol). The HEPES buffer (pH 7.2) was used as a matrix instead of a phosphate buffer to avoid the potential interaction of the metal(loid) and phosphate ions. The prepared tubes were incubated at room temperature for 5 min or overnight with continuous rotation to avoid sedimentation of the beads. After the incubation sample was immediately collected by Protocol A or Protocol B. In Protocol A, a filtrate not containing beads was collected using a spin column and used for the ICP quantification. In Protocol B, the beads were collected by centrifugation and subjected to the ICP analysis.

For Protocol B (Fig. 2), the reaction between Cd and the beads was examined. The whole reaction mixture (5 mL) was centrifuged at 4,255 g for 1 min immediately after the reaction. The supernatant was quickly but carefully removed by a vacuum aspirator not to disturb the bead pellet. The beads were resuspended with 1 mL of 60% nitric acid. The resuspension and another 1 ml of 60% nitric acid for rinse were transferred to a 10mL PTFE tube. The sample was digested in an open vessel at 140 °C for 3 h. The digested samples were filled up to 5 ml with 2% nitric acid, and their Cd concentrations were determined by ICP-OES. The quantity of Cd binding to control beads (M_[control beads]_) and to ligand-attached beads (M_[ligand beads]_) after the reaction was calculated, respectively (Fig. 2). Then, the amount of Cd interacted with the ligand attached to the beads (M_[binding]_) was obtained by subtracting M_[control beads]_ from M_[ligand beads]_ (Fig. 2).

## Results

### Preparation of phytochelatin- and glutathione-attached beads

PCs are commercially available only on a 1 mg scale (Eurogentec, Seraing, Belgium) as far as we searched, and it was a considerable cost disadvantage to our large demand of PCs for the affinity beads preparation. Thus, we first synthesized a shorter PC [(γ-Glu-Cys)_2_-Gly, PC2] and a longer PC [(γ-Glu-Cys)_3_-Gly, PC3] by a standard Fmoc solid-phase peptide synthesis protocol (Supplementary Fig. S1). The two different length PCs are common PC isoforms synthesized in *S. pombe*, *C. elegans,* and various plant cells exposed to toxic metal(loid) ions such as Cd(II) and As(III) [14,17,20,22,28]. Fmoc-Gly-OH was loaded onto 2-chlorotrityl chloride resin (1.38 mmol/g, 200 mg). Fmoc protecting group was removed using a solution of 2% 1,8-diazabicyclo[5.4.0]undec-7-ene (DBU) in *N,N*-dimethylformamide (DMF). Condensation of Fmoc-Cys(Trt)-OH and Fmoc-Glu-O*t*Bu was performed using hydroxybenzotriazole (HOBt) and *N,N’*-diisopropylcarbodiimide (DIPCI). The resulting peptide was cleaved from the resin with 20% 1,1,1,3,3,3-hexafluoro-2-propanol (HFIP) in CH_2_Cl_2_, and then deprotected with trifluoroacetic acid (TFA)/triisopropylsilane (TIPS)/H_2_O (95:2.5:2.5). After purification by preparative HPLC, PC2 and PC3 were obtained in 64% yield (95.0 mg) and 64% yield (126 mg), respectively. Then we analyzed the synthesized PC2 and PC3 by HPLC/MS to confirm the purity and exact mass. The synthesized PC2 was eluted as a peak with a retention time of 0.75 min in total ion current (TIC) chromatogram (Fig. 1A) and 0.68 min in photodiode array (PDA) chromatogram monitored at 210 nm (Fig. 1C) and was detected as a protonated molecule [M + H]+ at *m/z* 540.14 (Fig. 1E). The synthesized PC3 was eluted in a peak with a retention time of 1.68 min in TIC chromatogram (Fig. 1B) and 1.60 min in PDA chromatogram monitored at 210 nm (Fig. 1D) and was detected as a protonated molecule [M + H]+ at *m/z* 772.12 (Fig. 1F).

**Fig. 1.**
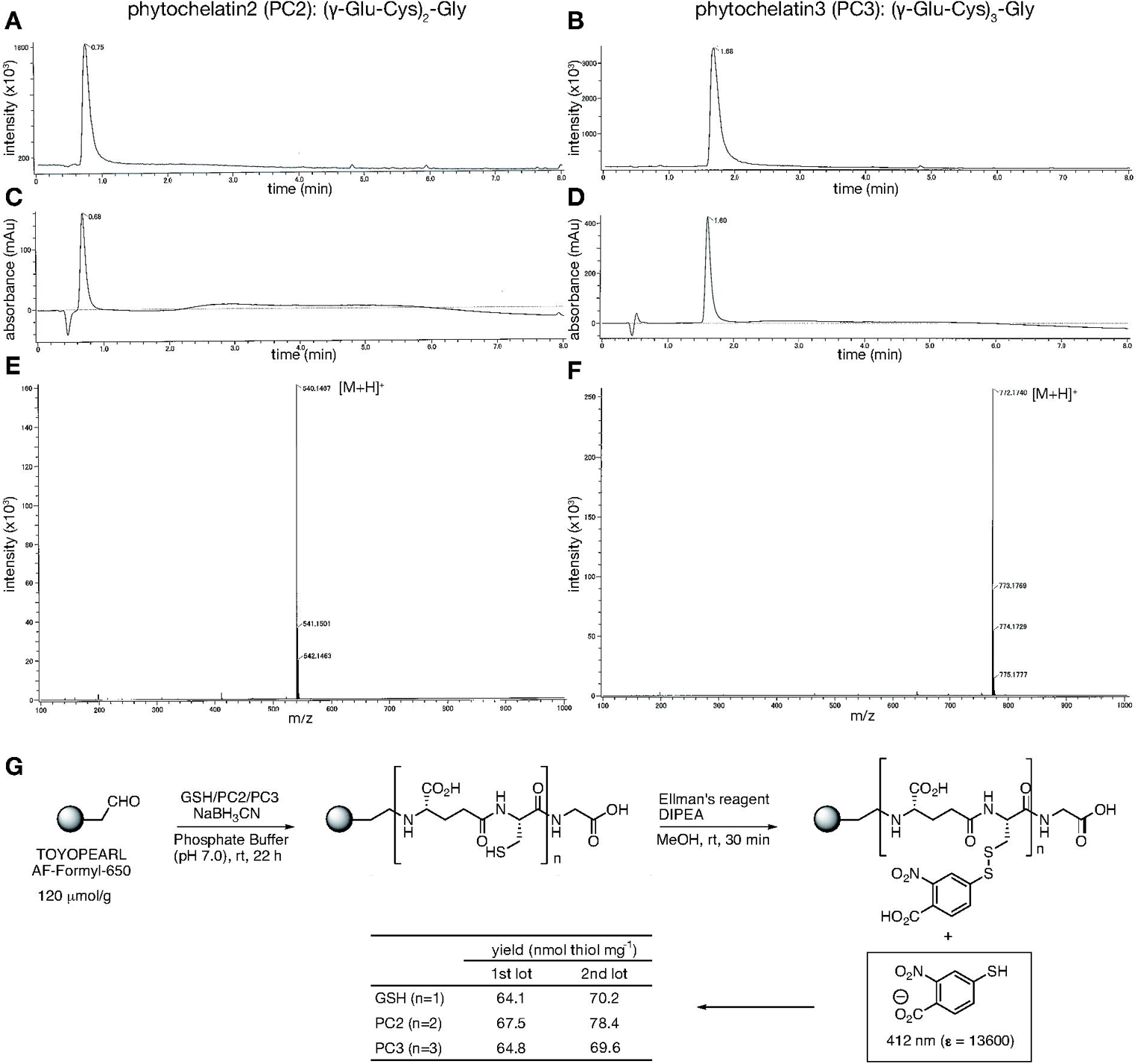
Verification of the synthesized phytochelatins. (A-F) and loading to the beads (G). (A, B) Total ion current chromatograms of the synthesized PC2 and PC3. (C, D) LC-PDA chromatograms of the synthesized PC2 and PC3 (at 210 nm). (E, F) Mass spectrum of the synthesized PC2 and PC3 (acquisition range: *m/z* 100–1000). (G) Loadings of GSH, PC2, and PC3 onto TOYOPEARL AF-Fornyl-650 beads were carried out under reductive amination conditions. The loadings were estimated by detecting the amount of 2-nitro-5-thiobenzoate anion released from the reaction of Ellman’s reagent and thiol groups on the affinity beads.

Using the synthetic PC2, PC3, and commercially obtained GSH, we attempted to prepare PC- or GSH-attached affinity beads (Fig. 1G). Hydrophilic beads which are only modified with formyl groups were used as a carrier resin. GSH, PC2, or PC3 was loaded onto the beads via reductive amination using NaBH_3_CN. This reaction was conducted twice independently to obtain two different preparations of the respective ligand-attached beads. The yield was quantified by Ellman’s reagent [29] and was overall within a similar range (64.1 to 78.4 nmol thiol per mg beads) irrespective of the ligands and preparations (Fig. 1G). The beads treated only with the reaction buffer not containing GSH, PC2, or PC3 were prepared as control beads in the subsequent binding assay.

### Establishment of the assay protocol for metal(loid)-ligand interaction

We then designed and tested protocols for evaluating the ligand-metal(loid) interactions using the prepared ligand-attached beads (Fig. 2). A reaction mixture for the standard protocol was prepared in a 5 ml plastic tube containing 3.0 mg of the beads and 200 nmol of metal(loid)-of-interest. 4-(2-hydroxyethyl)-1-piperazineethanesulfonic acid (HEPES) buffer (pH 7.2) was used as a matrix instead of a phosphate buffer to avoid the potential interaction of the metal(loid) and phosphate ions. The prepared tubes were incubated at room temperature for 5 min or overnight with continuous rotation to avoid sedimentation of the beads. Sample collection after the incubation was performed by two different protocols: collecting a filtrate without beads using a spin column (Protocol A) and collecting beads without a solution by centrifugation (Protocol B). Element contents in the collected samples were quantified by ICP-OES.

For calculation of [M]_binding_ [the metal(loid) binding to the ligand on the beads], in Protocol A, [M]_ligand filtrate_ [the metal(loid) amount in the filtrate from the ligand-attached beads reaction] was subtracted from [M]_control filtrate_ [the metal(loid) amount in the filtrate from the control beads reaction] (Fig. 2). In Protocol B, [M]_control beads_ [the metal(loid) amount in the control beads] was subtracted from [M]_ligand beads_ [the metal(loid) amount in the ligand-attached beads] to obtain [M]_binding_ (Fig. 2).

The two protocols were tested with Cd(II) as a model metal. To compare the accuracy of Protocol A and B, we first quantified total Cd adsorption to the control and ligand-attached beads (Fig. 3). For Protocol A, [Cd]_ligand filtrate_ and [Cd]_control filtrate_ were respectively subtracted from Cd(II) input to each reaction (200 nmol). For Protocol B, [Cd]_control beads_ and [Cd]_ligand beads_ were directly used to assess total Cd adsorption to the beads. Regardless of the types of the beads, the Cd adsorption to the beads did not significantly differ between Protocol A and B (Fig. 3), suggesting that both protocols can equally evaluate the metal binding to the beads as designed. Protocol A is more convenient and practical because the separation of the beads and the buffer can be easily and quickly achieved by a spin column centrifugation, whereas Protocol B requires careful but quick removal of the buffer by pipetting after centrifugation. Taken together, we decided to choose Protocol A for further experiments.

**Fig. 3.**
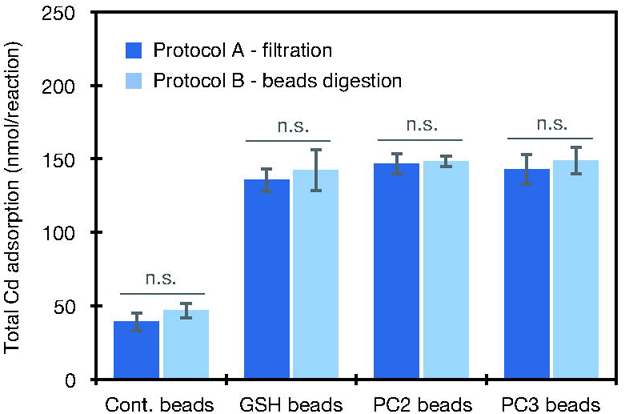
Comparison of Protocol A and Protocol B using Cd as a metal model. Total Cd adsorption to the respective types of affinity beads was analyzed by ICP-OES after the 5-min incubation with 200 nmol Cd. Data represent means with SD from three independent reactions for each combination. There is no significant difference between the two protocols for all affinity beads tested (*P* > 0.05, Student’s T-test). n.s., not significant.

### Affinity variation of GSH, PC2, and PC3 with various metal(loid)s

The experiment to test the two protocols using Cd(II) as a model metal indicated the substantial chelation between the ligands and Cd(II) because all the ligand-attached beads (GSH, PC2, and PC3) adsorbed nearly 3-times more Cd than the control beads (Fig. 3). The results motivated us to further conduct a full-scale experiment and we quantified the GSH and PCs-mediated chelation of Mn(II), Fe(II), Ni(II), Cu(II), Zn(II), As(III), and Cd(II) (Fig. 4). Note that As(III) does not have a charge at physiological pH. 5-min incubation was applied for the initial experiment, and overnight incubation was also examined for Mn(II), Fe(II), Cu(II), and As(III). The experiment for Fig. 4 was carried out with the first lot beads, except for Cd(II) for which the independently prepared second lot beads were also applied to examine possible lot-specific effects.

**Fig. 4.**
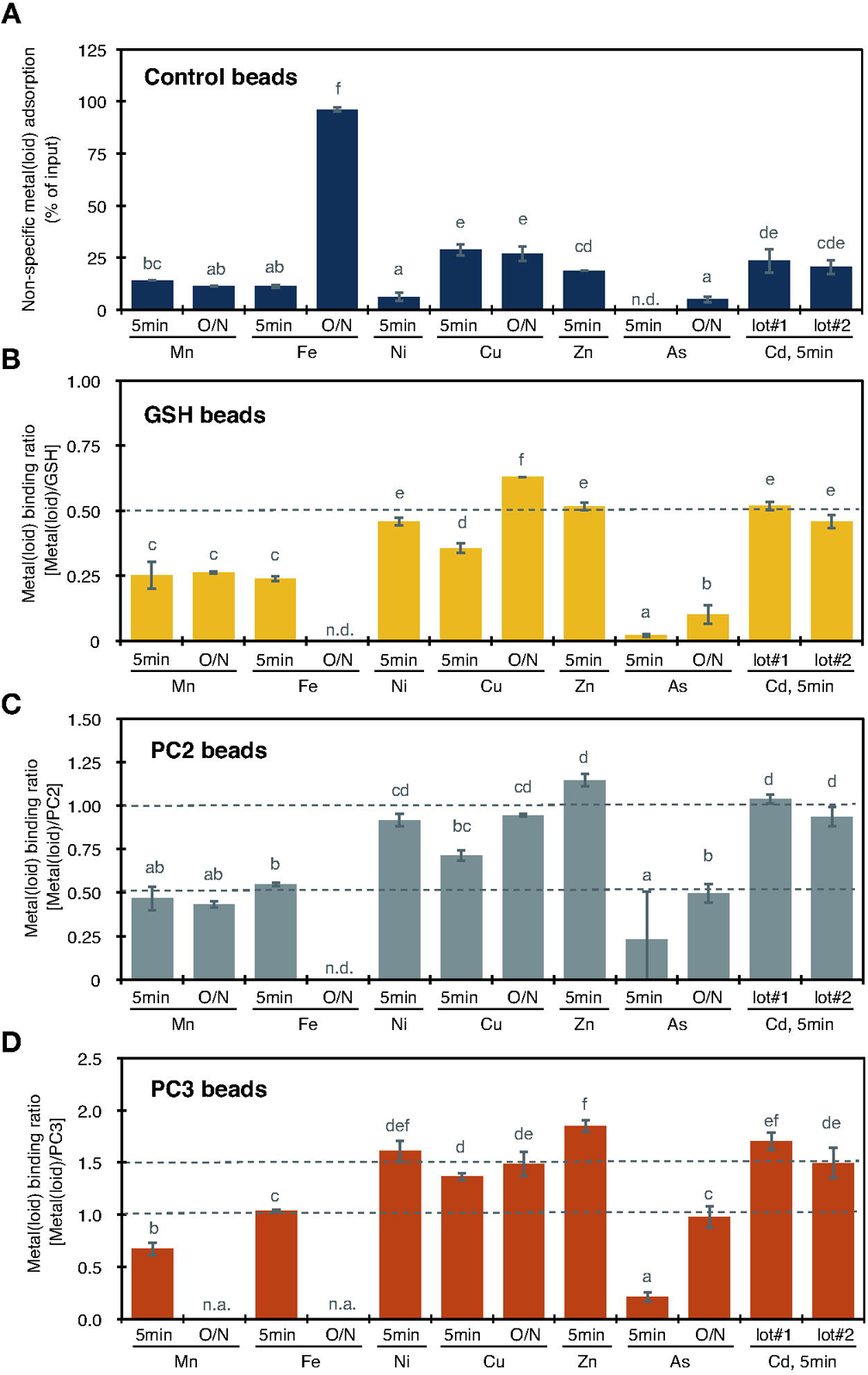
Metal(loid)-ligand interaction assay using the ligand-attached affinity beads. A reaction mixture containing 3.0 mg of each type of the beads and 200 nmol of each metal(loid) species was incubated for 5 min or overnight at room temperature with contentious rotation. After the incubation, the reaction mixture was immediately subjected to spin column filtration (Protocol A). The metal(loid) content in the filtrate was quantified by ICP-OES. Data represent means with SD from three independent reactions for each combination. Means sharing the same letter are not significantly different (*P* < 0.05, Tukey’s HSD). (A) Non-specific metal(loid) adsorption to the control (blank) beads. (B-D) Metal(loid) binding ratios of GSH-attached beads (B), PC2-attached beads (C), and PC3-attached beads (D). n.d., not detected. n.a., not analyzed.

We first checked the metal(loid) adsorption to the controls beads as non-specific background adsorption of the assay (Fig. 4A). The presented values are percentages of the metal(loid) binding to the beads to the total input to the reaction (200 nmol). Non-specific adsorption to the hydrophilic beads reached up to 30% of the input, regardless of the tested metal(loid) species and incubation time, except for overnight incubation with Fe(II) (nearly 100% of the input to the reaction was adsorbed to the carrier beads). The result of the non-specific background adsorption indicates that the experimental condition provides enough metal(loid)s left in the reaction buffer which can interact with the ligand on the ligand-attached beads, except for overnight Fe incubation.

Then we calculated the metal(loid) binding ratio of GSH (Fig. 4B), PC2 (Fig. 4C), and PC3 (Fig. 4D), dividing [M]_binding_ (Fig. 2) by the molar number of the ligand in the reaction. For GSH which harbors one cysteine, the metal-binding ratio of 0.5 was referred to as a standard to indicate a binding stoichiometry of one metal to two GSH. The results of 5-min incubation for Ni(II), Zn(II), and Cd(II) were close to 0.5 (Fig. 4B). The ratio of 5-min incubation with Cu(II) showed a somewhat intermediate value (0.36). Those of Mn(II) and Fe(II) were significantly lower even than this value of Cu(II), suggesting a lower affinity of Mn(II) and Fe(II) for GSH. For As(III), the binding ratio of 0.33 was referred to as a standard indicating a binding stoichiometry of one As(III) to three GSH [30]. However, the ratio of As/GSH after 5-min incubation was 0.02, which was far below the expected value of 0.33. To further examine the lower binding ratio of Cu(II), Mn(II), Fe(II), and As(III), the incubation time was prolonged from 5 min to overnight. The binding ratios of Mn(II) and As(II) with GSH after overnight incubation was still quite lower than the standards 0.5 and 0.33, respectively (Fig. 4B). The overnight incubation increased the binding ratio of Cu(II) and GSH from 0.36 (5-min) to 0.62, both of which were within the range of the standard value of 0.5. The binding of Fe(II) with GSH after the overnight incubation was not detected due to the huge background Fe adsorption to the carrier beads (Fig. 4A).

The metal(loid) binding ratio with PC2 is shown in Fig. 4C. For PC2, which harbors two cysteines, the binding ratios of 0.5 and 1.0 as standards were referred to indicate 1:2 and 1:1 binding of metal(loid) and PC2, respectively. The binding ratios of Ni(II), Cu(II) (overnight), Zn(II), and Cd(II) were statistically similar and were close to 1.0 (Fig. 4C). The ratio obtained from 5-min incubation with Cu(II) was again a bit lower than that from overnight incubation, but the difference was smaller than the case of GSH (Fig. 4B). Binding ratios with PC2 for Mn(II) (5-min and overnight) and Fe(II) (5-min) were far below 1.0. The ratio for As/PC2 was close to 0.5 after overnight incubation, suggesting 1:2 binding of As(III) and PC2, whereas the result of 5-min incubation was much lower.

The metal(loid) binding ratio with PC3 was shown in Fig. 4D. PC3 contains three cysteines, and the binding ratios of 1.0 and 1.5 as standards were referred to indicate 1:1 and 3:2 binding of metal(loid) and PC3, respectively. The binding ratios of Ni(II), Cu(II) (5-min and overnight), Zn(II), and Cd(II) were around 1.5. It should be noted that there was no statistical difference between the incubation periods for Cu(II). The ratios for Mn(II) (5-min) and Fe(II) (5-min) were 0.67 and 1.04, respectively, which were significantly lower than the ratios for other divalent metals tested. The binding ratio of As(III) and PC3 (overnight) was 0.98, indicating the 1:1 binding relationship, however, the 5-min incubation again resulted in a lower value (0.22).

### Effects of different beads preparation

Conducting the experiments to produce the most results for Fig. 3 and Fig. 4, we used a large portion of the first preparation of the ligand-attached beads. Thus, we prepared a second lot of the respective beads. The second preparation resulted in similar yields however, the yield was overall a bit higher (Fig. 1G). We examined the possible effects of the different lots on the metal(loid)-ligand-binding assay, using Cd(II) as a model again. Background non-specific Cd adsorption (Fig. 4A), and binding ratios with GSH (Fig. 4B), PC2 (Fig. 4C), and PC3 (Fig. 4D) were not statistically different between the first and second preparations by Tukey’s HSD. In addition, the results of the Student’s T-test were also not significant (*P* > 0.05). These results suggest that differently prepared ligand-attached beads can be equally used in the binding assay and we decided to use the second lot for further experiments.

### Binding competition between Cd and essential metals

The results of the single metal(loid)-ligand assay (Fig. 4) suggested different affinity of the small thiol compounds to the various essential metals and toxic metal(loid)s. We next planned to explore functional differences of the different length thiols, GSH, PC2, and PC3, using the developed ligand-attached beads system. PCs are believed to function in chelating free toxic metal(loid)s in the cytosol. However, competition over PCs’ binding sites can be expected in the cytosol between the toxic metals and co-existing essential heavy metals. Thus, we conducted a competition assay using Cd(II) as a model. Cd(II), not As(III), was selected because Cd(II) binding was stably detected by the 5-min incubation, whereas As(III) required the overnight incubation (Fig. 4). In this competition assay, Cd(II) (200 nmol) as well as essential metals [Mn(II) (20 nmol), Fe(II) (40 nmol), Cu(II) (2 nmol), and Zn(II) (20 nmol)] were added to the reaction. The ratio of the supplemented metals was derived from a rough calculation based on the total element concentrations in leaves of the plant model *Arabidopsis thaliana* [31–33].

As a result of the competition assay, Mn(II) and Fe(II) exhibited inferior affinity to GSH, PC2, and PC3 compared to Cu(II), Zn(II), and Cd (II) (Fig. 5A, B, C). Smaller fractions of the Mn(II) and Fe(II) input (~30%) were associated with the ligands, and approximately 60% or more of the Cu(II), Zn(II), and Cd(II) input were associated with the ligands. We then calculated profiles of the metals binding to GSH, PC2, and PC3, and compared them with the profile of the input (Fig. 5D). Reflecting the lower affinity with the ligands, the fractions of Mn(II) and Fe(II) binding to the ligands were smaller than those of the input. In contrast, the Cd(II) binding to the ligands was more than 80% of the total metals captured by the ligands, which was larger than the input supplemented Cd(II) fraction (73%). These results further support the results of the single-metal binding assay (Fig. 4), suggesting a preference of Cd(II) in binding to the thiol ligands, compared to the tested essential metals.

**Fig. 5.**
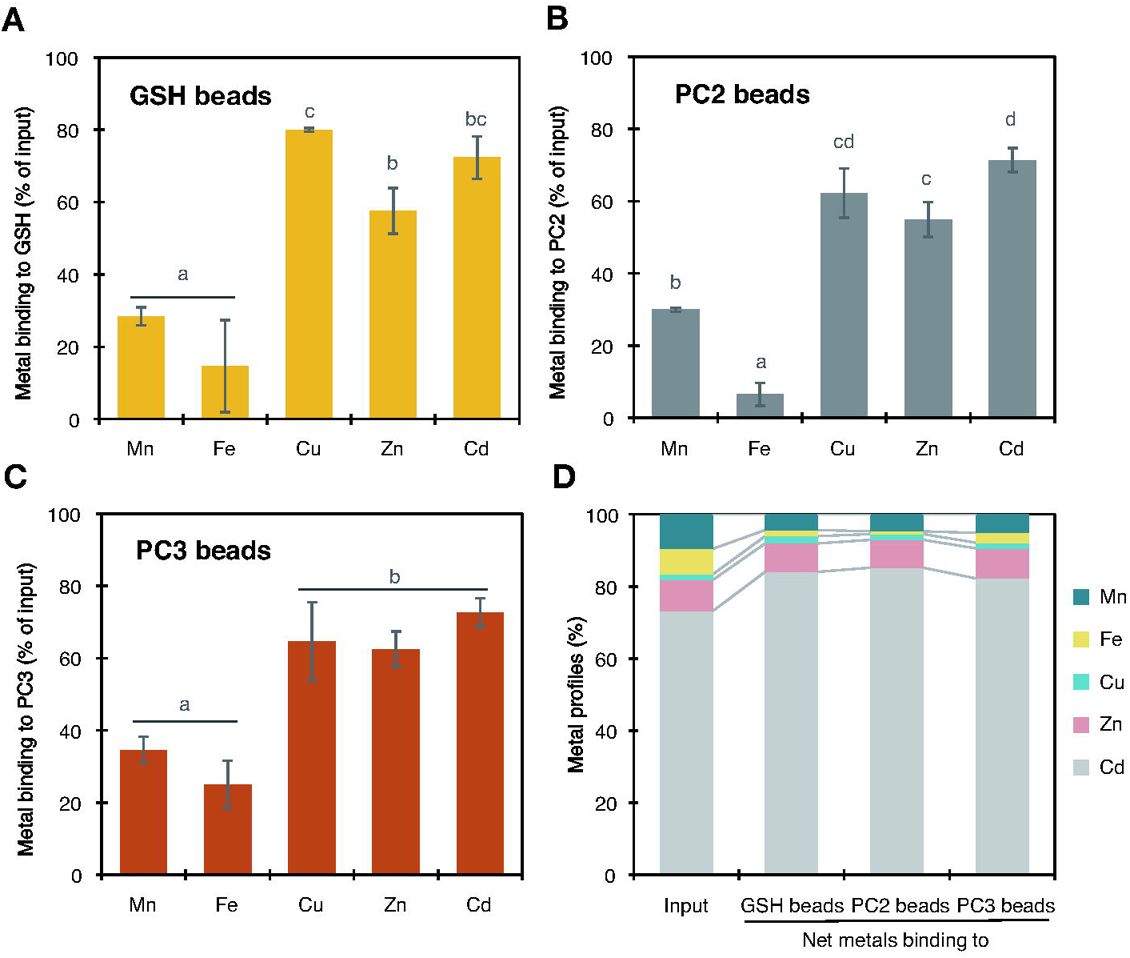
Metal competition assay using the ligand-attached beads. The reaction buffer containing 20 nmol of MnCl_2_, 40 nmol of FeCl_2_, 2 nmol of CuSO_4_, 20 nmol of ZnSO_4_, and 200 nmol of CdCl_2_ was added to 3.0 mg of each type of the beads. The tube was immediately set to a rotator and incubated at room temperature for 5 min. After the incubation, the reaction mixture was immediately subjected to spin column filtration (Protocol A). The metals content in the filtrate was quantified by ICP-OES. (A-C) Proportions of metals binding to GSH (A), PC2 (B), and PC3 (C) on the beads, calculated from total metal input to the reaction and the metals associated with each ligand on the beads. Data represent means with SD from three independent reactions. Means sharing the same letter are not significantly different (*P* < 0.05, Tukey’s HSD). (D) Profiles of the metals in the reaction mixture (input) and on the respective beads. Data represent means from three independent reactions.

### Cd-binding competition between GSH and PCs

Next, we conducted another competition assay to examine a possible preference of PC and Cd(II) interaction (Fig. 6). First, the binding of 100 nmol Cd(II) and GSH beads competed with or without 100 nmol PC2 added as a free form competitor in the reaction mixture. The Cd(II) binding to GSH beads under the condition without the competitor PC2 was 55% of the total Cd(II) input (Fig. 6A), but the addition of PC2 as the competitor to the reaction significantly decreased the Cd(II) binding to GSH beads to 16% of the total Cd(II) input (Fig. 6A). Then, the inverted assay conditions were also tested (Fig. 6B): Binding of 100 nmol Cd(II) by PC2 beads competed with 25 nmol GSH or 200 nmol GSH supplemented to the reaction mixture in a free form. In the absence of GSH, 55% of the input Cd(II) was bound with PC2 on the beads, and the addition of 25 nmol nor 200 nmol GSH to the mixture as a competitor did not change the Cd(II) binding profile to PC2 on the beads. The results overall suggest the superior binding affinity of Cd(II) and PCs compared to GSH.

**Fig. 6.**
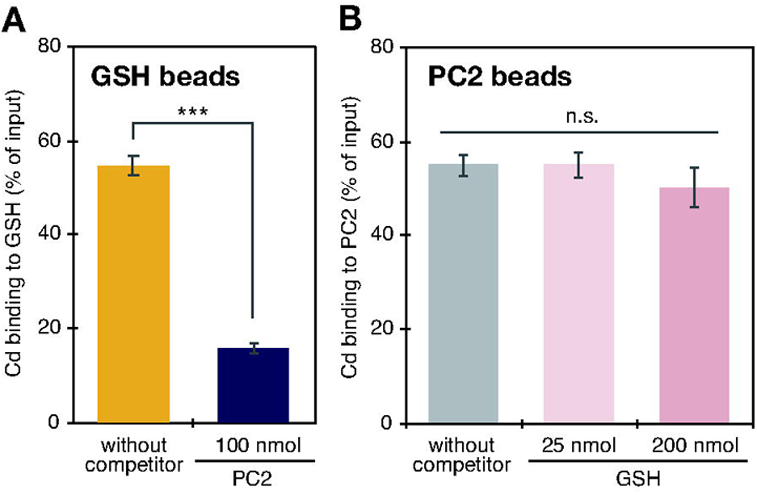
PC2 and GSH competition assay using the ligand-attached beads. (A) The reaction buffer containing 100 nmol of CdCl_2_ and 100 nmol of PC2 was added to 3.0 mg of the GSH-attached beads. (B) The reaction buffer containing 100 nmol of CdCl_2_, and 25 or 200 nmol of GSH was added to 3.0 mg of the PC2-attached beads. (A, B) The tube was immediately set to a rotator after adding the buffer and incubated at room temperature for 5 min. After the incubation, the reaction mixture was immediately subjected to spin column filtration (Protocol A). The Cd content in the filtrate was quantified by ICP-OES. Data represent means with SD from three independent reactions. *** *P* < 0.001, (Student’s T-test). n.s. *P* > 0.05 (Dunnett’s test).

## Discussion

### Advantages of our metal-ligand binding assay

Physiological roles of metal-ligand interactions have been attracting attention because of their significance in the essential metal homeostasis and toxic metal(loid) detoxification in bacteria, plants, and humans [26,34–36]. The first goal of this study was to establish a new assay system practically feasible for biologists interested in evaluating metal-biological ligand interactions under cellular-mimicking conditions. The idea of our assay resembles that of protein affinity purification protocols such as immunoprecipitation in biochemistry. Using ICP as a readout tool is the first unique advantage of our assay. ICP-OES and ICP-MS are widely used by metal biologists. Moreover, the application of ICP enables rapid, highly sensitive, and stable quantification of a large sample set. Note that a standard ICP protocol takes only 3-4 min for a single sample measurement of even more than 10 elements at ppb (ICP-OES) or sub-ppb levels (ICP-MS). In addition, a spin column-based efficient separation of the beads by centrifugation is also a key advantage of the assay. The spin-column procedure is very familiar to molecular biologists and enables immediate termination of the reaction of multiple samples. Besides technical advantages, it should be noted that our assay system allows variable assay conditions which mimic cellular environments. Thus, we can easily examine the effects of various co-existing metals and endogenous metal-interactants (Figs. 4-6). These features are unique to our assay and imply the broad utility of the protocol.

A few limitations exist in our assay: the assay cannot provide direct complex form information and stability constants unlike the other conventional methods such as potentiometry, voltammetry, NMR, ITC, and ESI-MS. However, the proposed assay has significant advantages in terms of flexibility of assay conditions, high-throughput, and technical feasibility especially for molecular biologists who are interested in metal-ligand interactions under cellular environments.

### GSH and metals interaction as a test model of the assay

In the present study, we employed GSH and two different length PCs (PC2 and PC3) as the model ligands for our assay (Figs. 1-3). First, the reliability of the assay was verified by assessing GSH-metal interactions (Fig. 4B). The affinity between GSH and various metals is well characterized by conventional analytical methods. GSH is believed to be the most likely ligands in the eukaryotic cells to form metal-complexes through its sulfhydryl group [8,10] (note that GSH analogs rather than GSH itself are prominent in prokaryotes [37]). The abundance in the cells is another advantage of GSH as a metal-ligand: GSH concentrations in eukaryotic cells (ranging from 0.1 mM to 8 mM [38–40]) are likely substantial enough to bind free metals which would exist at much lower concentrations in cells.

The results of binding experiments between GSH and well-studied cations [Mn(II), Fe(II), Ni(II), Cu(II), Zn(II), and Cd(II)] coincide with the reported results and the order of the Irving–Williams series (Mn<Fe<Co<Ni<Cu, Zn) [41] which describes the complex stability between divalent metal ions and organic ligands (note that Cd is not included in the series, and Co is not tested in this study). For example, our assay suggests a weak interaction of Mn(II) and Fe(II) with GSH (Fig. 4B), supporting the models suggested in the previous studies: Mn(II)-GSH and Fe(II)-GSH interactions are 1:1 with markedly lower stability constant (log K_1_ values) of 2.7 and 5.1, respectively [42,43]. In contrast, stronger affinity with GSH is suggested for highly reactive metals up in the Irving–Williams series [Ni(II), Cu(II), and Zn(II)] as well as Cd(II) (Fig. 4B). The reported stability constants (log β values) for interactions between GSH and these metals are around 10 or much higher even under physiological pH conditions [44,45]. In our study, the metal/GSH ratio of around 0.5 was obtained for Ni(II), Cu(II), Zn(II), and Cd(II) (Fig. 4B), suggesting 1:2 metal-GSH complex formation as previously reported [44,45]. For Cu(II), there would be other possibilities. The Cu/GSH ratios were slightly yet significantly lower than 0.5 under 5 min-incubating conditions, and overnight incubation increased the ratio (Fig. 4B). This variation would be explained by the controversial nature of Cu-GSH complex formation. Previous studies imply Cu(II) and GSH affinity with 1:2 and 1:1 stoichiometry [45] and in addition, the formation of Cu(I)_4_(GS)_6_ has been suggested [46]. It should be also noted that we applied Cu(II) to the assay, not Cu(I), because of its low solubility and extremely high background adsorption to the beads in the preliminary experiment using CuCl (data not shown). Cu(II) is converted to Cu(I) under the reducing cellular conditions [46], which may explain the increased interaction of Cu and GSH at the longer reaction period (Fig. 4B), probably caused by different stoichiometry between Cu(II)-GSH and Cu(I)-GSH. Therefore, taken together with these points, our results for Cu(II) cannot suggest an exact stoichiometry of Cu(II)-GSH interaction, but at least suggest a stronger affinity with GSH under physiological (cellular) conditions compared to Mn(II) and Fe(II) (Fig. 4B).

Nevertheless, the results of GSH beads (Fig. 4B) overall demonstrate that under physiological pH conditions, the assay can detect the different affinity of metals and GSH, corresponding to the order of the Irving–Williams series and the established stability constants.

### Preferential interaction between phytochelatin and Cd

Several metabolites other than GSH are also suggested as biological metal ligands. PC is an established example that mainly functions in Cd detoxification through Cd-PC complex formation in higher plants, *S. pombe*, and *C. elegans* [16,17,20,22]. Due to its significance in Cd detoxification, the PCS activation mechanism has been extensively examined using recombinant proteins, heterologous expression systems in *S. pombe*, plant cell cultures, and plant seedlings. In brief, PCS activity is triggered substantially by various environmentally relevant ions [e.g. As(III), As(V), Ag(I), Cd(II), Hg(II), and Pb(II)] and the excess of a few physiologically essential metal ions [Cu(II) and Zn(II)] as well, irrespective of their valency and positions in the periodic table [14,17–19,28,47–49]. On the other hand, several elements essential for plant nutrition [K(I), Mg(II), Ca(II), Mn(II), and Ni(II)] even in their excess levels do not activate PCSs [48,49]. Whether Fe(II) and Fe(III) activate PCSs is still controversial [49]. Such metal-(loid) PCS activation mechanism has been a significant question. Recent studies suggest that the different regions of the C-terminal regulatory domain of PCSs are required for the metal(loid)-specific activation [17,50–53], although the detailed enzymatic action of PCS remains elusive mainly due to the lack of the crystal structure of the full-length PCS.

In contrast to the activation mechanism, much less is known about the subsequent metal(loid)-PC complex formation. Only a few studies reported the detection of Cd-PC and As-PC complexes in plant extracts or *in vitro* samples using LC-ESI-MS or LC-ICP-MS [54–59]. Thus, the second purpose of the present study is to examine preferences of metal(loid)-PC complex formation using our binding assay, mainly focusing on Cd-PC interaction. The assay using PC2 beads and a single metal species (Fig. 4C) suggests a 1:1 interaction between Cd(II) and PC2. The 3:2 interaction between Cd(II) and PC3 is suggested in accordance with the three SH residues in PC3 (Fig. 4D). These interaction ratios are in line with the suggested values obtained by ESI-MS analysis of the plant samples and *in vitro* ITC analysis [57,60]. Similarly, our assay suggests 1:1 interaction with PC2 and 3:2 interaction with PC3 were also suggested for Ni(II), Cu(II), and Zn(II). Fe(II)-PC2 interaction is suggested weaker (Fig. 4C), but Fe(II) is likely to interact with PC3 at the ratio of 1:1 (Fig. 4D). These results suggest a potential of Fe(II), Ni(II), Cu(II), and Zn(II) interacting with PCs, aside from their PCS activation ability. This implies that Cd(II) and these metals may compete for the PCs’ binding sites. However, the Cd-PC complexes are the major form detected in plants exposed to Cd stress [56–58], suggesting that Cd(II)-PC interaction is dominant over the competitive essential metals.

A series of competitive assays were conducted to examine the dominance of Cd-PC interaction (Figs. 5-6). The competition assay between Cd(II) and several plant essential metals suggests the preferential binding affinity of PCs with Cd(II) over the tested essential metals (Fig. 5). The result supports the hypothesis and *in planta* observation of the dominance of Cd-PCs formation [56–58]. In plant cellular conditions, sub-millimolar levels of GSH exist as another potent Cd(II) chelator other than PCs. Thus, we examined the Cd(II) binding preference between PC2 and GSH (Fig. 6). The assays clearly demonstrate the marked preference of Cd-PC2 interaction over Cd-GSH (probably Cd-GS_2_). The Cd-PC2 interaction was not interfered with even by the excess level of free GSH (Fig. 6B), which supports the reported higher stability constant of Cd-PC2 than Cd-GS_2_ [45,61]. Overall, our competition assays suggest the robust and preferential interaction of PC and Cd(II) over other co-existing essential metals and GSH. It should be noted that the actual mechanism of PC-metal(loid) complex formation is still unclear. A previous study suggested a possibility that the pre-formed Cd-GS_2_ complex interacts with the C-terminal regulatory domain of the PCS and activates the transpeptidase activity to induce PC synthesis [62]. The previous study also proposed that a Cd ion is transferred from the Cd-GS_2_ complex to the PCS protein and activates PC synthesis [62]. Thus, it can be postulated that Cd-PC complex formation (partly) occurs during the process of PC synthesis [62]. Nevertheless, our competition assays (Figs. 5-6) suggest that the formed Cd-PC complex would be stable enough in the cytosol, not interfered by GSH and other essential metals. This stability would be significant to ensure subsequent transport of the Cd-PC complex from the cytosol to the vacuole, which is mediated by the tonoplast ABC transporters [63,64].

### Interaction between thiols and As(III)

Another promising partner of PC other than Cd(II) is As(III). In plants, As(III) stress also induces PC synthesis, and the subsequent As(III)-PC complex formation and vacuolar sequestration by ABC transporters are suggested to be crucial for As(III) detoxification [17,51,65,66]. Formation of As(III)-(PC2)_2_ and As(III)-PC3 is reported *in vitro* and *in planta* [54,55,59]. The binding stoichiometry is in accordance with the number of the thiol groups in PCs [note that three SH residues are required to catch one As(III)]. Our results of As(III)-PC beads interaction assay under the overnight condition clearly support these observations, with the As/PC2 ratio of 0.5 (As:PC2=1:2) and As/PC3 ratio of 1 (As:PC3=1:1) (Fig. 5C and 5D). In addition to PCs, As(III) would also interact with GSH and form As(GS)_3_ *in vitro* [67]. However, the complex *in vivo* is only observed under extreme biological conditions: the bile of rats injected with a very high dose of arsenite [68] and the roots of Arabidopsis PCS1 mutant *cad1-3* [59]. The instability of As(GS)_3_ at physiological pH and temperature [25] would partly explain why this species has only infrequently been observed *in vivo*. Supporting this hypothesis, the As/GSH ratio obtained under physiological conditions was much lower than 0.33 (Fig. 4B), the index for As(GS)_3_ formation. Taken together with the previous observations *in vitro* and *in vivo*, our results suggest that As(GS)_3_ complex is not a major form in the cells challenged by As(III) except for the extreme conditions, and an As-PC complex is much more stable [69] and thus crucial for the As detoxification in plants.

## Conclusion

The importance of toxic metals and essential metals homeostasis in plants as food resources is increasing to achieve better human health, and the small molecular weight ligands are major (potential) players in regulating these biological processes [26,70]. *In vitro* evaluation of metal-ligand interactions is a crucial initial step for characterizing ligand-based metal homeostasis in the cells. The proposed assay system in this study has advantages regarding throughput, sensitivity, robustness, and specificity. Another strong point is that the assay is flexible in terms of reaction conditions. Thus, it is suitable for testing various metal-ligand combinations. Taking all these advantages, the present study demonstrated dominant natures of PC-Cd interactions by a series of binding assays.

Beyond PCs, the assay can be applied to test the potent metal affinity of other ligands. For example, characterizing the metal interactions of the Fe/Zn-chelator NA or other siderophores under physiological matrix conditions would advance our understanding of the biological chelators. Other interesting ligand candidates include isoforms of GSH and PCs in which the C-terminal Gly is replaced by β-Ala, Ser, Gln, or Glu [71–73]. These iso-thiols are found in various plant species but have not been well characterized. From the viewpoint of environmental science/ecotoxicology, the assay can be used for the assessment of environmentally important metal(loid)s and biological ligands interaction. Recently, in addition to the “classical metals”, a number of elements are newly utilized by high-tech industries and their levels in the environment are accordingly increasing [74,75]. For these further applications, detection by ICP-MS instead of ICP-OES is an option that would increase the sensitivity of the assay and target elements to be tested. Another possible modification to the method includes the buffer (matrix) composition, depending on the application. For example, certain metalloproteins are known to adversely interact with HEPES buffer [76], suggesting that other buffer types should be considered for such cases.

## Abbreviations

DBU: 1,8-diazabicyclo[5.4.0]undec-7-ene
DIPCI: *N,N’*-diisopropylcarbodiimide
DMF: *N,N*-dimethylformamide
GSH: glutathione
HEPES: 4-(2-hydroxyethyl)-1-piperazineethanesulfonic acid
HFIP: 1,1,1,3,3,3-hexafluoro-2-propanol
HOBt: hydroxybenzotriazole
ICP-OES: inductively coupled plasma-optical emission spectrometry
ICT: isothermal titration calorimetry
LC-ESI-MS: liquid chromatography-electrospray ionization mass-spectrometry
LC-ICP-MS: liquid chromatography-inductively coupled plasma mass-spectrometry
NA: nicotianamine
NMR: nuclear magnetic resonance
PC: phytochelatin
PCS: phytochelatin synthase
TFA: trifluoroacetic acid
TIPS: triisopropylsilane.

## Acknowledgments

We thank Prof. Tohru Nagamitsu (Kitasato University) and Dr. Mitsuru Abo (Meiji University) for helpful advice and discussion. This work was partly supported by the Japan Society for the Promotion of Science (Grant nos. 18K05377, 21K05330 to SU).

## Author contributions

SU, KN, and MK designed the experiments and prepared the manuscript. SU, KN, FN, and YOt conducted the experiments. SU, KN, YOh, RN, YT, and MK analyzed the data. All authors discussed the results and approved the manuscript.

## Data availability

The data underlying this article are available in the article and in its online supplementary material.

## References

1. Waldron KJ, Rutherford JC, Ford D & Robinson NJ (2009) Metalloproteins and metal sensing. Nature 460, 823–30.

2. Andreini C, Bertini I, Cavallaro G, Holliday GL & Thornton JM (2008) Metal ions in biological catalysis: from enzyme databases to general principles. J Biol Inorg Chem 13, 1205–18.

3. Dupont CL, Butcher A, Valas RE, Bourne PE & Caetano-Anollés G (2010) History of biological metal utilization inferred through phylogenomic analysis of protein structures. Proc Natl Acad Sci USA 107, 10567–10572.

4. Dupont CL, Yang S, Palenik B & Bourne PE (2006) Modern proteomes contain putative imprints of ancient shifts in trace metal geochemistry. Proc Natl Acad Sci USA 103, 17822–17827.

5. Hitomi Y, Outten CE & O’Halloran TV (2001) Extreme zinc-binding thermodynamics of the metal sensor/regulator protein, ZntR. J Am Chem Soc 123, 8614–8615.

6. Outten CE & O’Halloran TV (2001) Femtomolar sensitivity of metalloregulatory proteins controlling zinc homeostasis. Science 292, 2488–2492.

7. Rae T, Schmidt P, Pufahl R, Culotta V & O’Halloran TV (1999) Undetectable intracellular free copper: The requirement of a copper chaperone for superoxide dismutase. Science 284, 805–808.

8. Colvin RA, Holmes WR, Fontaine CP & Maret W (2010) Cytosolic zinc buffering and muffling: their role in intracellular zinc homeostasis. Metallomics 2, 306–317.

9. Finney LA & O’Halloran TV (2003) Transition metal speciation in the cell: insights from the chemistry of metal ion receptors. Science 300, 931–936.

10. Hogstrand C, Kille P, Nicholson RI & Taylor KM (2009) Zinc transporters and cancer: a potential role for ZIP7 as a hub for tyrosine kinase activation. Trends Mol Med 15, 101–111.

11. Bashir K, Rasheed S, Kobayashi T, Seki M & Nishizawa NK (2016) Regulating subcellular metal Homeostasis: The key to crop improvement. Front Plant Sci 7, 1192.

12. Clemens S, Deinlein U, Ahmadi H, Höreth S & Uraguchi S (2013) Nicotianamine is a major player in plant Zn homeostasis. BioMetals 26, 623–632.

13. Ghssein G, Brutesco C, Ouerdane L, Fojcik C, Izaute A, Wang S, Hajjar C, Lobinski R, Lemaire D, Richaud P, Voulhoux R, Espaillat A, Cava F, Pignol D, Borezee-Durant E & Arnoux P (2016) Biosynthesis of a broad-spectrum nicotianamine-like metallophore in *Staphylococcus aureus*. Science 352, 1105–1109.

14. Grill E, Winnacker EL & Zenk MH (1987) Phytochelatins, a class of heavy-metal-binding peptides from plants, are functionally analogous to metallothioneins. Proc Natl Acad Sci USA 84, 439–443.

15. Howden R, Goldsbrough PB, Andersen CR & Cobbett CS (1995) Cadmium-sensitive, *cad1* mutants of *Arabidopsis thaliana* are phytochelatin deficient. Plant Physiol 107, 1059–1066.

16. Ha SB, Smith AP, Howden R, Dietrich WM, Bugg S, O’Connell MJ, Goldsbrough PB & Cobbett CS (1999) Phytochelatin synthase genes from Arabidopsis and the yeast *Schizosaccharomyces pombe*. Plant Cell 11, 1153–1164.

17. Uraguchi S, Tanaka N, Hofmann C, Abiko K, Ohkama-Ohtsu N, Weber M, Kamiya T, Sone Y, Nakamura R, Takanezawa Y, Kiyono M, Fujiwara T & Clemens S (2017) Phytochelatin synthase has contrasting effects on cadmium and arsenic accumulation in rice grains. Plant Cell Physiol 58, 1730–1742.

18. Fischer S, Kühnlenz T, Thieme M, Schmidt H & Clemens S (2014) Analysis of plant Pb tolerance at realistic submicromolar concentrations demonstrates the role of phytochelatin synthesis for Pb detoxification. Environ Sci Technol 48, 7552–7559.

19. Tennstedt P, Peisker D, Böttcher C, Trampczynska A & Clemens S (2009) Phytochelatin synthesis is essential for the detoxification of excess zinc and contributes significantly to the accumulation of zinc. Plant Physiol 149, 938–948.

20. Clemens S, Kim EJ, Neumann D & Schroeder JI (1999) Tolerance to toxic metals by a gene family of phytochelatin synthases from plants and yeast. EMBO J 18, 3325–3333.

21. Kondo N, Imai K, Isobe M, Goto T, Murasugi A, Wada-Nakagawa C & Hayashi Y (1984) Cadystin a and b, major unit peptides comprising cadmium binding peptides induced in a fission yeast ----- separation, revision of structures and synthesis. Tetrahedron Lett 25, 3869–3872.

22. Vatamaniuk OK, Bucher EA, Ward JT & Rea PA (2001) A new pathway for heavy metal detoxification in animals. Phytochelatin synthase is required for cadmium tolerance in *Caenorhabditis elegans*. J Biol Chem 276, 20817–20820.

23. Meister A (1991) Glutathione deficiency produced by inhibition of its synthesis, and its reversal; Applications in research and therapy. Pharmacol Therapeut 51, 155–194.

24. Wood BA & Feldmann J (2012) Quantification of phytochelatins and their metal(loid) complexes: critical assessment of current analytical methodology. Anal Bioanal Chem 402, 3299–3309.

25. Raab A, Meharg AA, Jaspars M, Genney DR & Feldmann J (2003) Arsenic–glutathione complexes—their stability in solution and during separation by different HPLC modes. J Anal Atom Spectrom 19, 183–190.

26. Clemens S (2019) Metal ligands in micronutrient acquisition and homeostasis. Plant Cell Environ 42, 2902–2912.

27. Fukuda T, Nagai K, Yagi A, Kobayashi K, Uchida R, Yasuhara T & Tomoda H (2019) Nectriatide, a Potentiator of Amphotericin B Activity from *Nectriaceae* sp. BF-0114. J Nat Prod 82, 2673–2681.

28. Cazalé AC & Clemens S (2001) *Arabidopsis thaliana* expresses a second functional phytochelatin synthase. FEBS Lett 507, 215–219.

29. Badyal JP, Cameron AM, Cameron NR, Coe DM, Cox R, Davis BG, Oates LJ, Oye G & Steel PG (2001) A simple method for the quantitative analysis of resin bound thiol groups. Tetrahedron Lett 42, 8531–8533.

30. Schmöger ME, Oven M & Grill E (2000) Detoxification of arsenic by phytochelatins in plants. Plant Physiol 122, 793–801.

31. Uraguchi S, Ohshiro Y, Otsuka Y, Tsukioka H, Yoneyama N, Sato H, Hirakawa M, Nakamura R, Takanezawa Y & Kiyono M (2020) Selection of agar reagents for medium solidification is a critical factor for metal(loid) sensitivity and ionomic profiles of *Arabidopsis thaliana*. Front Plant Sci 11, 503.

32. Verret F, Gravot A, Auroy P, Leonhardt N, David P, Nussaume L, Vavasseur A & Richaud P (2004) Overexpression of *AtHMA4* enhances root-to-shoot translocation of zinc and cadmium and plant metal tolerance. FEBS Lett 576, 306–312.

33. Semane B, Cuypers A, Smeets K, Belleghem FV, Horemans N, Schat H & Vangronsveld J (2007) Cadmium responses in *Arabidopsis thaliana*: glutathione metabolism and antioxidative defence system. Physiol Plant 129, 519–528.

34. Li Y, Nguyen M, Baudoin M, Vendier L, Liu Y, Robert A & Meunier B (2019) Why is tetradentate coordination essential for potential copper homeostasis regulators in Alzheimer’s Disease? Eur J Inorg Chem 2019, 4712–4718.

35. Fasae KD, Abolaji AO, Faloye TR, Odunsi AY, Oyetayo BO, Enya JI, Rotimi JA, Akinyemi RO, Whitworth AJ & Aschner M (2021) Metallobiology and therapeutic chelation of biometals (copper, zinc and iron) in Alzheimer’s disease: Limitations, and current and future perspectives. J Trace Elem Med Bio 67, 126779.

36. Kenney GE, Dassama LMK, Pandelia ME, Gizzi AS, Martinie RJ, Gao P, DeHart CJ, Schachner LF, Skinner OS, Ro SY, Zhu X, Sadek M, Thomas PM, Almo SC, Bollinger JM, Krebs C, Kelleher NL & Rosenzweig AC (2018) The biosynthesis of methanobactin. Science 359, 1411–1416.

37. Fahey RC (2013) Glutathione analogs in prokaryotes. Biochim Biophys Acta Gen Subj 1830, 3182–3198.

38. Soboll S, Gründel S, Harris J, Kolb-Bachofen V, Ketterer B & Sies H (1995) The content of glutathione and glutathione S-transferases and the glutathione peroxidase activity in rat liver nuclei determined by a non-aqueous technique of cell fractionation. Biochem J 311, 889–894.

39. Sun X, Shih AY, Johannssen HC, Erb H, Li P & Murphy TH (2006) Two-photon imaging of glutathione levels in intact brain indicates enhanced redox buffering in developing neurons and cells at the cerebrospinal fluid and blood-brain interface. J Biol Chem 281, 17420–17431.

40. Gutiérrez-Alcalá G, Gotor C, Meyer AJ, Fricker M, Vega JM & Romero LC (2000) Glutathione biosynthesis in Arabidopsis trichome cells. Proc Natl Acad Sci 97, 11108–11113.

41. Irving H & Williams RJP (1953) The stability of transition-metal complexes. J Chem Soc 3, 3192–3210.

42. Martin RB & Edsall JT (1959) The association of divalent cations with glutathione 1. J Am Chem Soc 81, 4044–4047.

43. Hider RC & Kong XL (2011) Glutathione: a key component of the cytoplasmic labile iron pool. Biometals 24, 1179–1187.

44. Krężel A & Bal W (2004) Studies of zinc(II) and nickel(II) complexes of GSH, GSSG and their analogs shed more light on their biological relevance. Bioinorg Chem Appl 2, 293–305.

45. Walsh MJ & Ahner BA (2013) Determination of stability constants of Cu(I), Cd(II) & Zn(II) complexes with thiols using fluorescent probes. J Inorg Biochem 128, 112–123.

46. Morgan MT, Nguyen LAH, Hancock HL & Fahrni CJ (2017) Glutathione limits aquacopper(I) to sub-femtomolar concentrations through cooperative assembly of a tetranuclear cluster. J Biol Chem 292, 21558–21567.

47. Maitani T, Kubota H, Sato K & Yamada T (1996) The composition of metals bound to class III metallothionein (phytochelatin and its desglycyl peptide) induced by various metals in root cultures of *Rubia tinctorum*. Plant Physiol 110, 1145–1150.

48. Oven M, Page JE, Zenk MH & Kutchan TM (2002) Molecular characterization of the homo-phytochelatin synthase of soybean *Glycine max*: relation to phytochelatin synthase. J Biol Chem 277, 4747–4754.

49. Loscos J, Naya L, Ramos J, Clemente MR, Matamoros MA & Becana M (2006) A reassessment of substrate specificity and activation of phytochelatin synthases from model plants by physiologically relevant metals. Plant Physiol 140, 1213–1221.

50. Li M, Barbaro E, Bellini E, Saba A, Toppi LS di & Varotto C (2020) Ancestral function of the phytochelatin synthase C-terminal domain in inhibition of heavy metal-mediated enzyme overactivation. J Exp Bot 71, 6655–6669.

51. Uraguchi S, Sone Y, Ohta Y, Ohkama-Ohtsu N, Hofmann C, Hess N, Nakamura R, Takanezawa Y, Clemens S & Kiyono M (2018) Identification of C-terminal regions in *Arabidopsis thaliana* phytochelatin synthase 1 specifically involved in activation by Arsenite. Plant Cell Physiol 59, 500–509.

52. Kühnlenz T, Hofmann C, Uraguchi S, Schmidt H, Schempp S, Weber M, Lahner B, Salt DE & Clemens S (2016) Phytochelatin synthesis promotes leaf Zn accumulation of *Arabidopsis thaliana* plants grown in soil with adequate Zn supply and is essential for survival on Zn-contaminated soil. Plant Cell Physiol 57, 2342–2352.

53. Ruotolo R, Peracchi A, Bolchi A, Infusini G, Amoresano A & Ottonello S (2004) Domain organization of phytochelatin synthase: functional properties of truncated enzyme species identified by limited proteolysis. J Biol Chem 279, 14686–14693.

54. Raab A, Feldmann J & Meharg AA (2004) The nature of arsenic-phytochelatin complexes in *Holcus lanatus* and *Pteris cretica*. Plant Physiol 134, 1113–1122.

55. Bluemlein K, Raab A, Meharg AA, Charnock JM & Feldmann J (2008) Can we trust mass spectrometry for determination of arsenic peptides in plants: comparison of LC–ICP–MS and LC–ES-MS/ICP–MS with XANES/EXAFS in analysis of *Thunbergia alata*. Anal Bioanal Chem 390, 1739–1751.

56. Sadi BBM, Vonderheide AP, Gong J-M, Schroeder JI, Shann JR & Caruso JA (2008) An HPLC-ICP-MS technique for determination of cadmium–phytochelatins in genetically modified *Arabidopsis thaliana*. J Chromatogr B 861, 123–129.

57. Yen T, Villa JA & DeWitt JG (1999) Analysis of phytochelatin–cadmium complexes from plant tissue culture using nano electrospray ionization tandem mass spectrometry and capillary liquid chromatography/electrospray ionization tandem mass spectrometry. J Mass Spectrom 34, 930–941.

58. Chen L, Guo Y, Yang L & Wang Q (2007) SEC-ICP-MS and ESI-MS/MS for analyzing *in vitro* and *in vivo* Cd-phytochelatin complexes in a Cd-hyperaccumulator *Brassica chinensis*. J Anal At Spectrom 22, 1403–1408.

59. Liu W-JJ, Wood BA, Raab A, McGrath SP, Zhao F-JJ & Feldmann J (2010) Complexation of arsenite with phytochelatins reduces arsenite efflux and translocation from roots to shoots in Arabidopsis. Plant Physiol 152, 2211–2221.

60. Jacquart A, Brayner R, Chahine J-MEH & Ha-Duong N-T (2017) Cd^2+^ and Pb^2+^ complexation by glutathione and the phytochelatins. Chem-biol Interact 267, 2–10.

61. Dorčák V & Krężel A (2003) Correlation of acid–base chemistry of phytochelatin PC2 with its coordination properties towards the toxic metal ion Cd(II). Dalton Trans, 2253–2259.

62. Vatamaniuk OK, Mari S, Lang A, Chalasani S, Demkiv LO & Rea PA (2004) Phytochelatin synthase, a dipeptidyltransferase that undergoes multisite acylation with gamma-glutamylcysteine during catalysis: stoichiometric and site-directed mutagenic analysis of *Arabidopsis thaliana* PCS1-catalyzed phytochelatin synthesis. J Biol Chem 279, 22449–22460.

63. Mendoza-Cózatl DG, Zhai Z, Jobe TO, Akmakjian GZ, Song W-YY, Limbo O, Russell MR, Kozlovskyy VI, Martinoia E, Vatamaniuk OK, Russell P & Schroeder JI (2010) Tonoplast-localized Abc2 Transporter Mediates Phytochelatin Accumulation in Vacuoles and Confers Cadmium Tolerance. J Biol Chem 285, 40416–40426.

64. Park J, Song W-YY, Ko D, Eom Y, Hansen TH, Schiller M, Lee TG, Martinoia E & Lee Y (2012) The phytochelatin transporters AtABCC1 and AtABCC2 mediate tolerance to cadmium and mercury. Plant J 69, 278–288.

65. Song W-YY, Yamaki T, Yamaji N, Ko D, Jung K-HH, Fujii-Kashino M, An G, Martinoia E, Lee Y & Ma JF (2014) A rice ABC transporter, OsABCC1, reduces arsenic accumulation in the grain. Proc Natl Acad Sci 111, 15699–15704.

66. Song W-YY, Park J, Mendoza-Cózatl DG, Suter-Grotemeyer M, Shim D, Hörtensteiner S, Geisler M, Weder B, Rea PA, Rentsch D, Schroeder JI, Lee Y & Martinoia E (2010) Arsenic tolerance in Arabidopsis is mediated by two ABCC-type phytochelatin transporters. Proc Natl Acad Sci 107, 21187–21192.

67. Rey NA, Howarth OW & Pereira-Maia EC (2004) Equilibrium characterization of the As(III)–cysteine and the As(III)–glutathione systems in aqueous solution. J Inorg Biochem 98, 1151–1159.

68. Kala SV, Neely MW, Kala G, Prater CI, Atwood DW, Rice JS & Lieberman MW (2000) The MRP2/cMOAT transporter and arsenic-glutathione complex formation are required for biliary excretion of arsenic. J Biol Chem 275, 33404–33408.

69. Bluemlein K, Raab A & Feldmann J (2009) Stability of arsenic peptides in plant extracts: off-line versus on-line parallel elemental and molecular mass spectrometric detection for liquid chromatographic separation. Anal Bioanal Chem 393, 357–366.

70. Clemens S (2019) Safer food through plant science: reducing toxic element accumulation in crops. J Exp Bot 70, 5537–5557.

71. Klapheck S, Fliegner W & Zimmer I (1994) Hydroxymethyl-phytochelatins [γ-glutamylcysteine)n-serine] are metal-induced peptides of the *Poaceae*. Plant Physiol 104, 1325–1332.

72. Klapheck S, Schlunz S & Bergmann L (1995) Synthesis of phytochelatins and homo-phytochelatins in *Pisum sativum* L. Plant Physiol 107, 515–521.

73. Meuwly P, Thibault P, Schwan AL & Rauser WE (1995) Three families of thiol peptides are induced by cadmium in maize. Plant J 7, 391–400.

74. Merschel G & Bau M (2015) Rare earth elements in the aragonitic shell of freshwater mussel *Corbicula fluminea* and the bioavailability of anthropogenic lanthanum, samarium and gadolinium in river water. Sci Total Environ 533, 91–101.

75. Tokumaru T, Ozaki H, Onwona-Agyeman S, Ofosu-Anim J & Watanabe I (2017) Determination of the extent of trace metals pollution in soils, sediments and human hair at e-waste recycling site in Ghana. Arch Environ Contam Toxicol 73, 377–390.

76. Jahromi EZ, White W, Wu Q, Yamdagni R & Gailer J (2010) Remarkable effect of mobile phase buffer on the SEC-ICP-AES derived Cu, Fe and Zn-metalloproteome pattern of rabbit blood plasma. Metallomics 2, 460–468.

