## Supporting information for "Development of affinity beads-based *in vitro* metal-ligand binding assay reveals dominant cadmium affinity of thiol-rich small peptides phytochelatins beyond glutathione"

**Materials and chemicals**

All reagents for the synthesis were purchased at the highest commercial quality and used without further purification unless specified otherwise. TOYOPEARL AF-Formyl-650 was purchased from TOSOH (Tokyo, Japan).

**Loading of Fmoc-Gly-OH onto the 2-Chlorotrityl resin**

2-Chlorotrityl chloride resin (200 mg, 1.38 mmol/g, 100–200 mesh, 1% divinylbenzene polystyrene copolymer) was placed in a polypropylene tube and swollen in $CH_2Cl_2$ (8 mL) for 30 min. To the resin was added a solution of Fmoc-Gly-OH (280 mg, 960 μmol) and *N*,*N*-diisopropylethylamine (328 μL, 1.92 mmol) in $CH_2Cl_2$/DMF (4:1, v/v, 8 mL). The resulting mixture was agitated at room temperature for 3 h. The reaction was quenched with MeOH (0.5 mL), and stirring continued further for 10 min. The solvents and soluble reagents were removed by filtration, and the resin was washed with $CH_2Cl_2$ (8 mL, 1 min × 3), DMF (8 mL, 1 min × 3), and $CH_2Cl_2$ (8 mL, 1 min × 3).

**Coupling of Fmoc-Cys(Trt)-OH (Method A)**

The Fmoc-Gly-resin was placed in a polypropylene tube. The resin was treated with 2% 1,8-diazabicyclo[5.4.0]undec-7-ene (DBU) in DMF (8 mL) at room temperature for 1 min. The solvents and soluble reagents were removed by filtration, and the resin was washed with $CH_2Cl_2$ (8 mL, 1 min × 3), DMF (8 mL, 1 min × 3), and $CH_2Cl_2$ (8 mL, 1 min × 3). To this washed resin was added a solution of Fmoc-Cys(Trt)-OH (564 mg, 960 μmol) and hydroxybenzotriazole (HOBt) (142 mg, 1.05 mmol) in $CH_2Cl_2$/DMF (4:1, v/v, 8 mL), and then DIPCI (148 μL, 960 μmol) was added. The resulting mixture was agitated at room temperature for 3 h. The solvents and soluble reagents were removed by filtration, and the resin was washed with $CH_2Cl_2$ (8 mL, 1 min × 3), DMF (8 mL, 1 min × 3), and $CH_2Cl_2$ (8 mL, 1 min × 3).

**Coupling of Fmoc-Glu-OtBu (Method B)**

The Fmoc-Cys(Trt)-Gly-resin was placed in a polypropylene tube and swollen in DMF (8 mL). The resin was treated with 2% DBU in DMF (8 mL) at room temperature for 1 min. The solvents and soluble reagents were removed by filtration, and the resin was washed with $CH_2Cl_2$ (8 mL, 1 min × 3), DMF (8 mL, 1 min × 3), and $CH_2Cl_2$ (8 mL, 1 min × 3). To the peptidyl resin was added a solution of Fmoc-Glu-O*t*Bu (408 mg, 960 μmol) and HOBt (142 mg, 1.05 mmol) in $CH_2Cl_2$/DMF (4:1, v/v, 8 mL), and then DIPCI (148 μL, 960 μmol) was added. The resulting mixture was agitated at room temperature for 3 h. The solvents and soluble reagents were removed by filtration, and the resin was washed with $CH_2Cl_2$ (8 mL, 1 min × 3), DMF (8 mL, 1 min × 3), and $CH_2Cl_2$ (8 mL, 1 min × 3). Methods A and B were repeated twice for phytochelatin 2 (PC2) and 3 times for phytochelatin 3 (PC3).

**Cleavage from the resin**

The Fmoc-[γ-Glu(α-O*t*Bu)-Cys(Trt)]$_2$-Gly-resin was placed in a polypropylene tube. The resin was treated with 2% DBU in DMF (8 mL) at room temperature for 1 min. The solvents and soluble reagents were removed by filtration, and the resin was washed with $CH_2Cl_2$ (8 mL, 1 min × 3), DMF (8 mL, 1 min × 3), and $CH_2Cl_2$ (8 mL, 1 min × 3). The washed resin was treated with 20% 1,1,1,3,3,3-hexafluoro-2-propanol (HFIP) in $CH_2Cl_2$ (8 mL) at room temperature for 30 min. The resin was filtered and washed with 10% $CH_3OH$ in $CH_2Cl_2$ (8 mL, 1 min × 3). The collected filtrates were evaporated to afford the crude protected peptide, which was used for the next reaction without purification.

**Deprotection**

The crude protected peptide was treated with trifluoroacetic acid/triisopropylsilane/$H_2O$ (95.0/2.5/2.5, 10 mL) and dithiothreitol (150 mg, 960 μmol) at room temperature for 4 h. After removal of the solvent, the residue was purified by preparative HPLC to give the desired peptide. **H-[γ-Glu-Cys]$_2$-Gly-OH (PC2)**: Yield 64% (12 steps, 95.0 mg, white foamy solid), HPLC retention time 0.68 min, purity 99%, ESI-MS: calculated for $C_{18}H_{30}N_5O_{10}S_2$ $[M + H]^+$ 540.1434, found *m/z* 540.1425. **H-[γ-Glu-Cys]$_3$-Gly-OH (PC3)**. Yield 64% (16 steps, 126 mg, white foamy solid), HPLC retention time 1.56 min, purity 99%, ESI-MS: calculated for $C_{26}H_{42}N_7O_{14}S_3$ $[M + H]^+$ 772.1951, found *m/z* 772.1944.

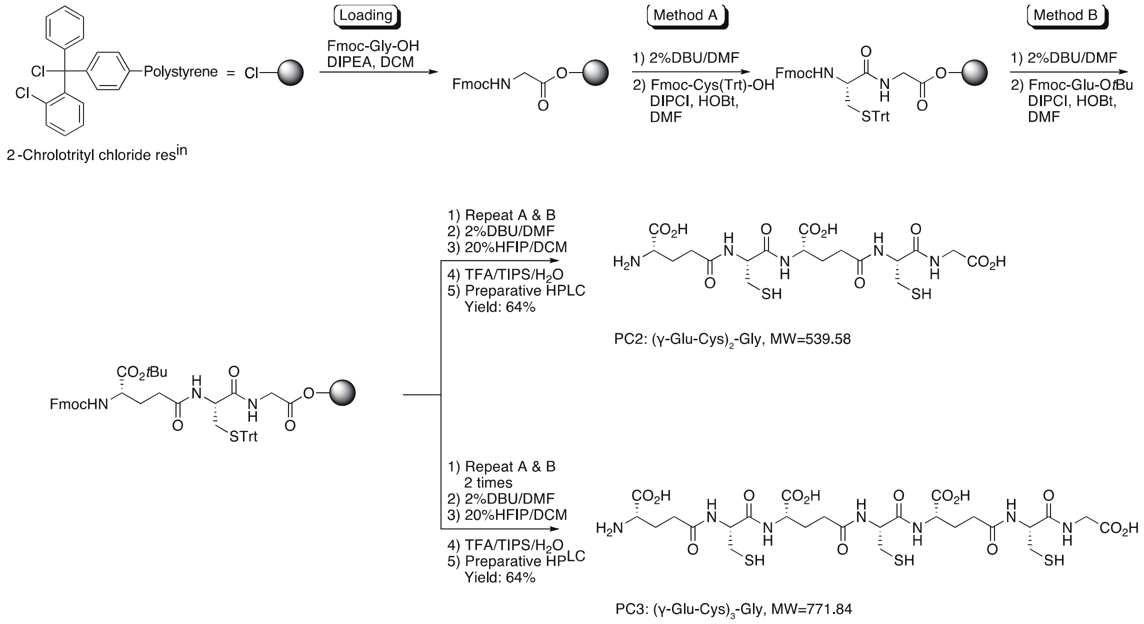


**Supplemental Figure S1** Synthesis of phytochelatins 2 and 3 (PC2 and PC3) by Fmoc solid-phase peptide synthesis using a 2-chlorotrityl chloride resin.
